# Measuring the long arm of childhood in real-time: Epigenetic predictors of BMI and social determinants of health across childhood and adolescence

**DOI:** 10.1101/2023.01.20.524709

**Authors:** Laurel Raffington, Lisa Schneper, Travis Mallard, Jonah Fisher, Liza Vinnik, Kelseanna Hollis-Hansen, Daniel A. Notterman, Elliot M. Tucker-Drob, Colter Mitchell, Kathryn P. Harden

## Abstract

Children who are socioeconomically disadvantaged are at increased risk for high body mass index (BMI) and multiple diseases in adulthood. The developmental origins of health and disease hypothesis proposes that early life conditions affect later-life health in a manner that is only partially modifiable by later-life experiences. Epigenetic mechanisms may regulate the influence of early life conditions on later life health. Recent epigenetic studies of adult blood samples have identified DNA-methylation sites associated with higher BMI and worse health (epigenetic-BMI).

Here, we used longitudinal and twin study designs to examine whether epigenetic predictors of BMI developed in adults are valid biomarkers of child BMI and are sensitive to early life social determinants of health. Salivary epigenetic-BMI was calculated from two samples: (1) N=1,183 8-to-19-year-olds (609 female, *mean* age=13.4) from the Texas Twin Project (TTP), and (2) N=2,020 children (1,011 female) measured at 9 and 15 years from the Future of Families and Child Well-Being Study (FFCWS).

We found that salivary epigenetic-BMI is robustly associated with children’s BMI (*r*=0.36 to *r*=0.50). Longitudinal analysis suggested that epigenetic-BMI is highly stable across adolescence, but remains both a leading and lagging indicator of BMI change. Twin analyses showed that epigenetic-BMI captures differences in BMI between monozygotic twins. Moreover, children from more disadvantaged socioeconomic status (SES) and marginalized race/ethnic groups had higher epigenetic-BMI, even when controlling for concurrent BMI, pubertal development, and tobacco exposure. SES at birth relative to concurrent SES best predicted epigenetic-BMI in childhood and adolescence.

We show for the first time that epigenetic predictors of BMI calculated from pediatric saliva samples are valid biomarkers of childhood BMI that are sensitive to social inequalities. Our findings are in line with the hypothesis that early life conditions are especially important factors in epigenetic regulation of later life health. Research showing that health later in life is linked to early life conditions have important implications for the development of early-life interventions that could significantly extend healthy life span.

---

Children who are socioeconomically disadvantaged are at increased risk for high body mass index (BMI) and multiple diseases in adulthood (Biro & Wien, 2010). The developmental origins of health and disease hypothesis proposes that early life conditions affect later-life health in a manner that is only partially modifiable by later-life experiences (Barker, 1990; Hayward & Gorman, 2004). In-utero and early postnatal development have received the most research attention as sensitive periods. Adolescence is another important developmental period that establishes foundations for adult health (Kankaanpää et al., 2022; Sawyer et al., 2012).

Epigenetic mechanisms, including DNA methylation, are thought to be involved in the biological embedding of early life conditions that affects aging-related health (Aristizabal et al., 2019; Watowich, 2022). Next-generation DNA-methylation predictors of biological aging that were developed to predict multi-system physiological decline, health behaviors, and/or mortality are promising new tools to study social determinants of health (Raffington & Belsky, 2022). A closely related set of studies has developed epigenetic predictors of BMI based on analysis of adult blood samples (epigenetic-BMI) (Hamilton et al., 2019; McCartney et al., 2018; Wahl et al., 2017). Similar to measures of biological aging, epigenetic-BMI has been found to improve the forecasting of health and mortality, including levels of triglycerides, HbA1c, HDL cholesterol, type 2 diabetes and cardiovascular disease, beyond phenotypic BMI and chronological age in adults (Hamilton et al., 2019; McCartney et al., 2018). It is likely that somewhat similar exposures and biological processes affect epigenetic predictors of biological aging and epigenetic-BMI, because (1) metabolic processes appear to be causally involved in biological aging and (2) next-generation epigenetic measures of biological aging are developed to predict BMI among other measures (Belsky et al., 2022; Hu et al., 2021).

Nascent studies in pediatric saliva samples suggest that epigenetic profiles of biological aging developed in adults are sensitive to social determinants of health experienced in real-time during childhood and adolescence (Niccodemi, 2022; Raffington et al., 2020, 2022). These studies suggest that salivary epigenetic predictors developed to predict adult health may be new tools to (1) study the early-life social determinants of lifelong health and (2) assess the efficacy of childhood interventions and policies by providing new surrogate outcomes relevant for health throughout the lifespan. Compared to blood collections, saliva samples are especially amenable to pediatric and hard-to-reach samples.

However, there remains considerable uncertainty about the validity of applying epigenetic predictors that were developed in adult blood samples to children’s saliva samples for three reasons. First, DNA-methylation is a primary mechanism of cell differentiation and is therefore tissue specific (Bakulski et al., 2016). Second, tissue composition across the body varies with age, which could confound comparisons across the lifespan (St-Onge & Gallagher, 2010). Third, genomic research has overwhelmingly relied on analysis of blood samples from adults of European ancestry, limiting the portability of DNAm-measures to diverse samples of children, for whom saliva is the most common and feasible biofluid to collect.

A challenge to validating epigenetic predictors of biological aging in children is that there is no gold standard measure of biological aging, especially early in the life course when people are generally healthy. In contrast, examining associations of salivary epigenetic-BMI with child BMI presents a unique opportunity to potentially validate the usage of blood-based DNA-methylation measures developed to predict adult health in pediatric saliva samples.

Here, we examine (1) whether epigenetic-BMI previously developed in adult blood samples is a valid biomarker of children’s BMI, when measured in saliva DNA-methylation, and (2) whether child epigenetic-BMI is sensitive to social determinants of health, as indexed by socioeconomic status and marginalized racial and ethnic identities in childhood and adolescence. To accomplish these goals, we analyze data from two demographically diverse pediatric cohorts that combine longitudinal and twin study designs. Our research builds on a previous blood-based epigenome-wide study that identified 278 CpG sites associated with BMI in 5387 adults of European and/or Indian Asian ancestry (Wahl et al., 2017). We use these results to compute salivary epigenetic-BMI in N=1,183 8-to 18-year-olds from the Texas Twin Project (TTP) and in N=2,020 children measured at ages 9 and again at 15 years in the Future of Families and Child Well-Being Study (FFCW).

## Methods

### Sample

*The Texas Twins Project* is an ongoing longitudinal study that includes the collection of saliva samples for DNA and DNA-methylation extraction since 2012 (Harden et al., 2013). Participants in the current study were 1213 (622 female) children and adolescents, including 433 monozygotic and 780 dizygotic twins from 617 unique families, aged 8 to 19 years (*M*=13.66, *SD*=3.06), who had at least one DNA-methylation sample. 195 participants contributed two DNA-methylation samples (time between repeated samples: *M*=22 months, *SD*=6.5, range 3 to 38 months), and 16 samples were assayed in duplicate for reliability analyses (total methylation sample n=1424). Participants self-identified as White only (*n*=752, 62%), Latinx only (*n*=147, 12%), Latinx and White (*n*=97, 8%), Black and potentially another race/ethnicity (*n*=120, Black, 10%), Asian and potentially another race/ethnicity but not Latinx or Black (*n*=90, Asian, 7.5%), and Indigenous American, Pacific Islander or other, but not Latinx, Black, or Asian (*n*=7, 0.6%). The University of Texas Institutional Review board granted ethical approval.

*The Future of Families and Child Wellbeing Study* (FFCW) follows a sample of 4,898 children born in large US cities during 1998-2000. FFCW oversampled children born to unmarried parents. During home visits BMI was measured, and saliva DNA was collected at ages 9 and 15 (N=3,100). Saliva DNA-methylation data were assayed on a 2/3 sample (n=2,020) using the Illumina 450K and EPIC methylation arrays with ages 9 and 15 assayed on the same plate. DNA-methylation study participants self-identified as Black/African-American only (*n*=1,009, 50%), Latinx (*n*=444, 22%), White (*n*=399, 20%), Multiracial (n=116, 5%), Asian/Other (*n*=52, 2%).

See **Table 1** for description of study measures, Supplemental Methods for preprocessing of DNA and DNA-methylation, and **Table S1** for descriptive statistics.

**Table 1.**
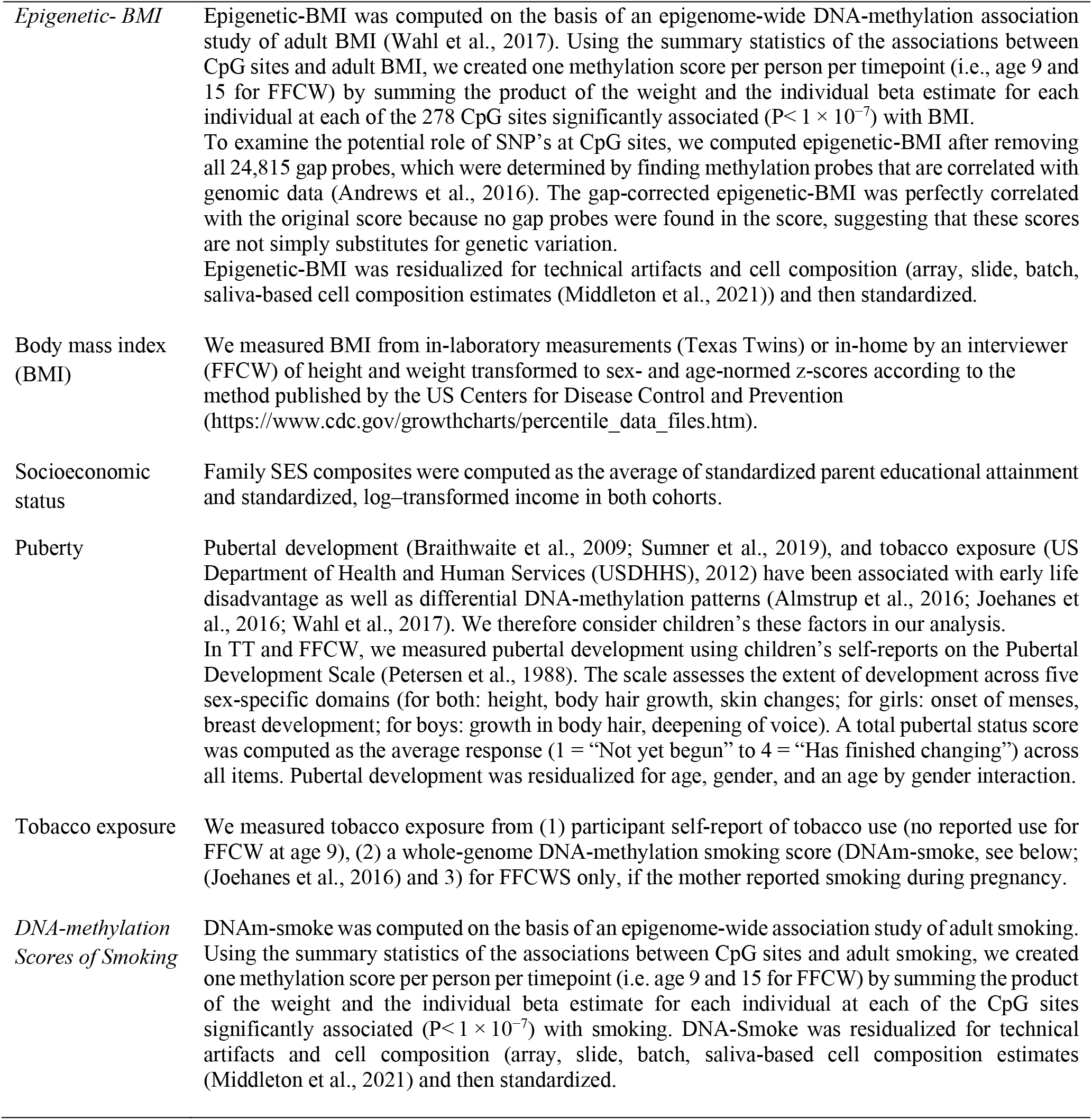
Description of study measures.

|  |  |
| --- | --- |
| *Epigenetic- BMI* | Epigenetic-BMI was computed on the basis of an epigenome-wide DNA-methylation association study of adult BMI (Wahl et al., 2017). Using the summary statistics of the associations between CpG sites and adult BMI, we created one methylation score per person per timepoint (i.e., age 9 and 15 for FFCW) by summing the product of the weight and the individual beta estimate for each individual at each of the 278 CpG sites significantly associated ($P < 1 \times 10^{-7}$) with BMI.<br>To examine the potential role of SNP's at CpG sites, we computed epigenetic-BMI after removing all 24,815 gap probes, which were determined by finding methylation probes that are correlated with genomic data (Andrews et al., 2016). The gap-corrected epigenetic-BMI was perfectly correlated with the original score because no gap probes were found in the score, suggesting that these scores are not simply substitutes for genetic variation.<br>Epigenetic-BMI was residualized for technical artifacts and cell composition (array, slide, batch, saliva-based cell composition estimates (Middleton et al., 2021)) and then standardized. |
| Body mass index (BMI) | We measured BMI from in-laboratory measurements (Texas Twins) or in-home by an interviewer (FFCW) of height and weight transformed to sex- and age-normed z-scores according to the method published by the US Centers for Disease Control and Prevention (https://www.cdc.gov/growthcharts/percentile_data_files.htm). |
| Socioeconomic status | Family SES composites were computed as the average of standardized parent educational attainment and standardized, log-transformed income in both cohorts. |
| Puberty | Pubertal development (Braithwaite et al., 2009; Sumner et al., 2019), and tobacco exposure (US Department of Health and Human Services (USDHHS), 2012) have been associated with early life disadvantage as well as differential DNA-methylation patterns (Almstrup et al., 2016; Joehanes et al., 2016; Wahl et al., 2017). We therefore consider children's these factors in our analysis.<br>In TT and FFCW, we measured pubertal development using children's self-reports on the Pubertal Development Scale (Petersen et al., 1988). The scale assesses the extent of development across five sex-specific domains (for both: height, body hair growth, skin changes; for girls: onset of menses, breast development; for boys: growth in body hair, deepening of voice). A total pubertal status score was computed as the average response (1 = "Not yet begun" to 4 = "Has finished changing") across all items. Pubertal development was residualized for age, gender, and an age by gender interaction. |
| Tobacco exposure | We measured tobacco exposure from (1) participant self-report of tobacco use (no reported use for FFCW at age 9), (2) a whole-genome DNA-methylation smoking score (DNAm-smoke, see below; (Joehanes et al., 2016) and 3) for FFCWS only, if the mother reported smoking during pregnancy. |
| *DNA-methylation Scores of Smoking* | DNAm-smoke was computed on the basis of an epigenome-wide association study of adult smoking. Using the summary statistics of the associations between CpG sites and adult smoking, we created one methylation score per person per timepoint (i.e. age 9 and 15 for FFCW) by summing the product of the weight and the individual beta estimate for each individual at each of the CpG sites significantly associated ($P < 1 \times 10^{-7}$) with smoking. DNA-Smoke was residualized for technical artifacts and cell composition (array, slide, batch, saliva-based cell composition estimates (Middleton et al., 2021) and then standardized. |

## Results

We conducted six sets of analyses. Results from analyses 1-3 validated the use of salivary epigenetic-BMI in children as a biomarker for BMI. Results from analyses 4-6 showed that epigenetic-BMI measures in children were sensitive to social determinants of health.

Prior to all analyses, epigenetic-BMI scores were residualized for technical artifacts, including array (Illumina Epic or 450k chips), slide, batch and estimated salivary cell composition. Analyses of duplicate samples suggested moderate reliability of epigenetic-BMI profiles (FFCW: 216 replicates ICC=0.67; TTP: 15 replicates ICC=0.43). All models included age, sex, and an age by sex interaction as covariates of epigenetic-BMI. Older children had higher epigenetic-BMI (*i.e.*, higher scores indicate increased risk of larger BMI score; TTP*: r*=0.33, CI=0.20 to 0.46, *p*<0.001; FFCW: mean difference between 9- and 15-year-olds=1.06, CI=0.60 to 1.54, *p*<0.001). Boys had lower epigenetic-BMI than girls (TTP: *b*=-0.11, CI=-0.20 to −0.02, *p*=0.018; FFCW-age9: *b*=-0.08, CI=-0.14 to 0.02, *p*=0.07; FFCW-age15: *b*=-0.24, CI=-0.30 to −0.18, *p*=0.008).

### (1) BMI gradients are reproduced in children’s salivary epigenetic-BMI

First, we tested if BMI was concurrently associated with epigenetic-BMI in both samples. In a multiple regression model where BMI *z*-scores were regressed on epigenetic-BMI (**Figure 1A**), epigenetic-BMI was significantly associated with BMI in 8- to 18-year-olds from the TTP (*r*=0.50, CI=0.42 to 0.59, *p*<0.001), FFCW 9-year-olds (*r*=0.36, 95% CI=0.31 to 0.40, *p*<0.001), and FFCW 15-year-olds (*r*=0.41, CI=0.36 to 0.45, *p*<0.001). The association between epigenetic-BMI and BMI was not accounted for by race/ethnicity, pubertal development, or indices of tobacco exposure (self-reported tobacco use in the TTP, maternal report of smoking during pregnancy in the FFCW, and DNA-methylation indicator of tobacco exposure in both samples), which were included as statistical controls (**Supplementary Table S2**). Compared to a model with only these covariates, the Δ*R*^2^ for epigenetic-BMI was 24.3% in TTP, 9.5% in FFCW-age9 and 13.6% in FFCW-age15.

**Figure 1.**
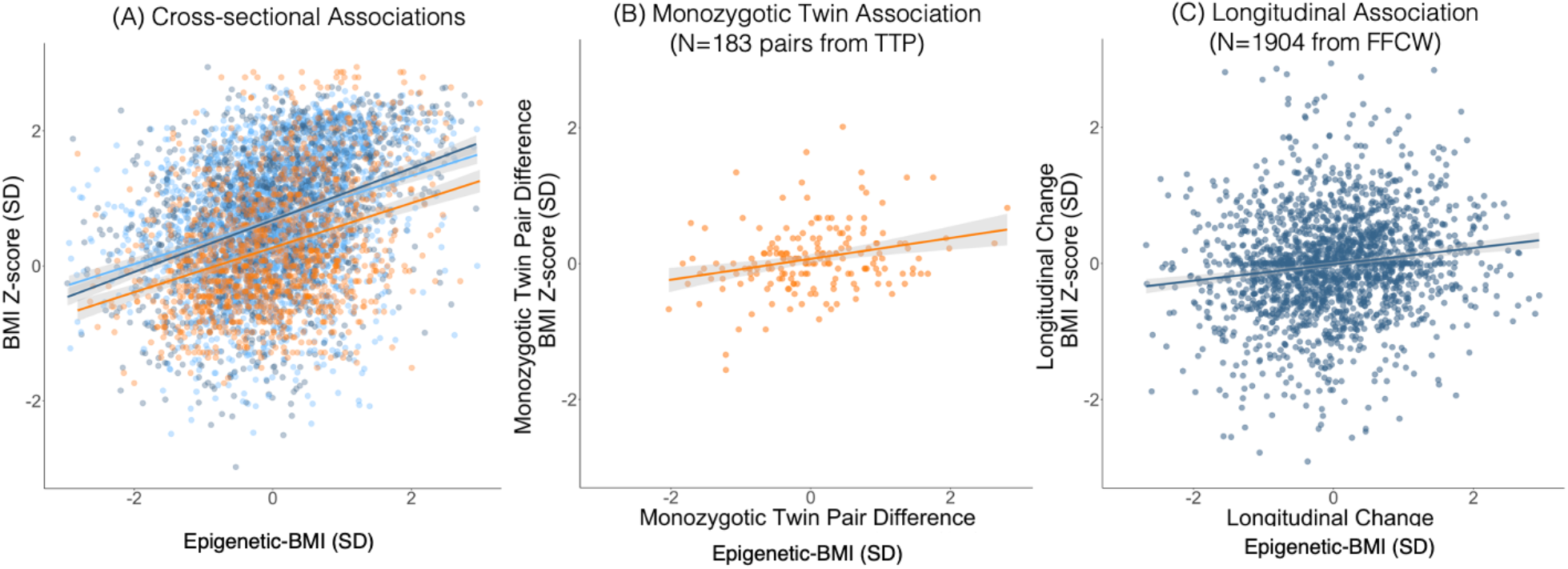
**(A) Cross-sectional associations between scaled epigenetic-BMI and measured BMI.** Results are presented for three samples: 8- to 18-year-old children from the Texas Twin Project (TTP), 9-year-old children and 15-year-old children from Future of Families and Child Wellbeing Study (FFCW-9; FFCW-15). **Epigenetic-BMI** and BMI z-scores were scaled in the full sample of each study and timepoint. **(B) Within monozygotic twin pair associations between scaled epigenetic-BMI and measured BMI.** Results based on N = 183 monozygotic twin pairs from TTP. (C) **Association of within-person longitudinal changes in scaled epigenetic-BMI and within-person change in BMI from age 9 to age 5**. Results based on N = 1904 longitudinal observations from FFCW.

**Figure 2.**
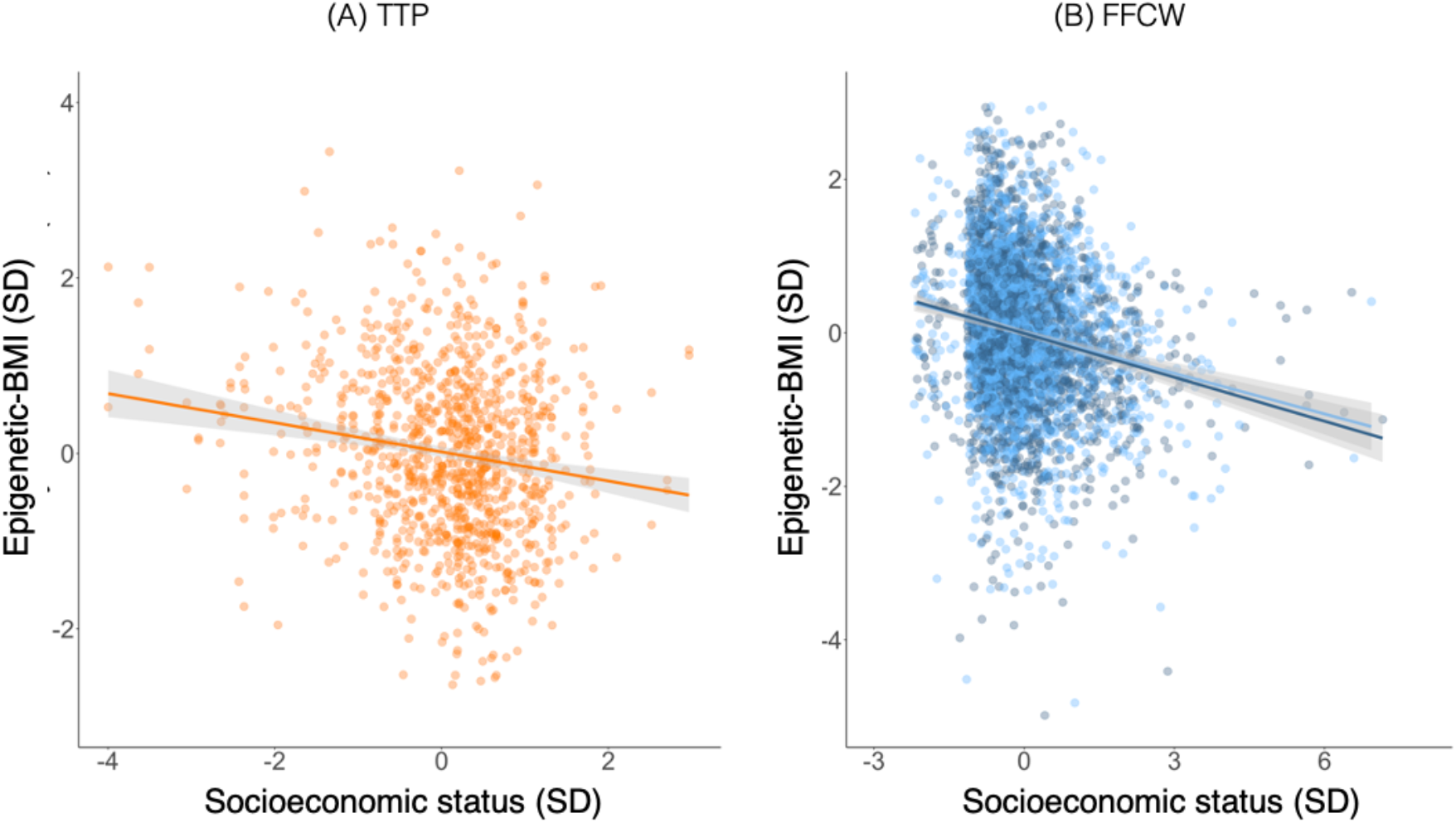
Socioeconomic inequalities in children’s epigenetic-BMI profiles. Crosssectional associations between scaled family-level socioeconomic status and salivary epigenetic-BMI. Results are presented for three samples: Panel A shows data form 8- to 18-year-old children from the Texas Twin Project (TTP), panel B shows data from 9-year-old children and 15-year-old children from Future of Families and Child Wellbeing Study (FFCW-9; FFCW-15). Socioeconomic status z-scores and epigenetic-BMI were scaled in the full sample of each study and timepoint.

### (2) Longitudinal analysis finds that epigenetic-BMI is stable across adolescence but still tracks changes in BMI

Prior analyses of cross-sectional data do not inform if epigenetic-BMI is a stable or highly variable measure across adolescence. Epigenetic-BMI measured at age 9 was strongly correlated with epigenetic-BMI measured at age 15 (b=0.63, CI=0.60 to 0.67, *p*<0.001).

Next, using longitudinal data from the FFCW, we fit a fixed-effects regression model to examine the correlation of within-person changes in BMI with changes in epigenetic-BMI from age 9 to 15 years, adjusting for unobserved time-invariant variables (**Figure 1C**; **Supplementary Table S3**). (In data like the FFCW, with exactly two observations per person, the person fixed-effects model reduces to an OLS regression using observed difference scores for the time-varying outcome variable). Within-person variation in epigenetic-BMI from ages 9 to 15 was associated with greater longitudinal increases in phenotypic BMI (*b*=0.13, CI=0.07 to 0.20, *p*<0.001), even after adjusting for longitudinal changes in SES and pubertal development. Epigenetic-BMI in the fixed-effects model explained 16% of the between-person variance in BMI but 2% of the variance in within-person change. As a negative control, within-person epigenetic-BMI from ages 9 to 15 was unassociated with changes in phenotypic height (*b*=0.00, CI=-0.8 to 0.09, *p*=0.87).

Next, we fit a bivariate random intercept cross-lagged panel model to test whether BMI at age 9 predicted epigenetic-BMI scores at age 15 and vice versa (Hamaker et al., 2015). We found evidence of a bi-directional relationship between epigenetic-BMI and BMI (**Supplementary Table S4, Supplemental Figure S2**): age-9 BMI predicted age-15 epigenetic-BMI (b=0.11, CI=.08 to 0.14, *p* <0.001) above and beyond age-9 epigenetic-BMI, and, in reverse, age-9 epigenetic-BMI predicted age-15 BMI (b=0.06, CI=.03 to 0.10, *p*<0.001), above and beyond age-9 BMI (note that, because stable variation in both DNAm and BMI is controlled for, the magnitude of the epigenetic-BMI → future BMI association is attenuated relative to the cross-sectional association). These results provide valuable evidence that salivary epigenetic-BMI can incrementally improve forecasting of future body size, over and above the child’s observed phenotype, even accounting for time-invariant factors such as stable genetic effects.

### (3) Epigenetic-BMI reflects differences in BMI between monozygotic twins

Epigenetic mechanisms, including DNA-methylation, are partially heritable (Hannon et al., 2018). To test the extent to which epigenetic-BMI was sensitive to exposures and developmental processes that make monozygotic twins different from each other, we used a monozygotic co-twin control analysis in the TTP. Monozygotic (*i.e*., identical) twins are nearly identical for their DNA sequence, so if differences in epigenetic-BMI between twins are associated with differences between them in their phenotypic BMI, this association is unlikely to be attributable to genes that influence both epigenetic-BMI and BMI.

This analysis was parameterized as a bivariate biometric model (*i.e.*, “ACE” model) that used the covariation between monozygotic twins and dizygotic twins to decompose the association between epigenetic-BMI and BMI into components representing additive genetic factors (A), environmental factors shared by twins living in the same home (C), and environmental factors unique to each twin (E).

Consistent with previous work showing that DNA-methylation is influenced by genetic variation, the heritability of epigenetic-BMI was estimated to be 46% (CI=21% to 71%), and the genetic correlation between epigenetic-BMI and measured BMI was moderate (rA=0.33, CI=0.17 to 0.50). However, the correlation between the E components of variation in DNAm and phenotypic BMI (rE), which reflects the extent to which identical twins who differ from their co-twins in epigenetic-BMI show corresponding differences in their BMI, was also positive and significant (**Supplementary Table S5;** see **Figure 1B**; rE=0.24, CI=0.10 to 0.37, *p*<0.001). Thus, among individuals who have been matched on nuclear DNA sequence, as well as on the background environmental factors shared by twins raised in the same family, variation in epigenetic-BMI continues to be associated with BMI. The moderate heritability of epigenetic-BMI indicates that, as with phenotypic BMI, there were both genetic and environmental sources of variation in methylation. The positive genetic and nonshared environmental correlations between epigenetic-BMI and phenotypic BMI indicate that it is both these genetic and environmental sources of variation in epigenetic and phenotypic BMI that are linked, which is consistent with a causal basis for the observed association.

Additionally, consistent with results from previous studies in adults (Reed et al., 2020; Shah et al., 2015; Trejo Banos et al., 2020), we found that salivary epigenetic-BMI provides complementary information compared to measured genetic variants associated with BMI (*i.e.*, polygenic indices of BMI, See **Supplementary Results** and **Table S6**).

### (4) Children from lower socioeconomic status homes have higher epigenetic-BMI

In the next set of analyses, we tested if epigenetic-BMI is sensitive to social determinants of health. First, we tested if children growing up in more socioeconomically disadvantaged circumstances exhibited higher concurrent epigenetic-BMI. Epigenetic-BMI was significantly predicted by family-level socioeconomic status (SES), measured using composites of parental income and education (Error! Reference source not found.**; Supplementary Table S7**). Results from the TTP sample were as follows: *b*=-0.24, CI=-0.34 to −0.13, *p*<0.001. Results from the FFCW age 9: *b*=-0.17, CI=-0.22 to −0.13, *p*<0.001; FFCW age 15: *b*=-0.19, CI=-0.23 to −0.14, *p*<0.001.

We next examined whether the association between socioeconomic disadvantage and epigenetic-BMI was robust to controlling for concurrent BMI, pubertal development and tobacco exposure (**Supplementary Table S7).** In both cohorts, even among children of comparable body size, pubertal development, and tobacco exposure, children from lower SES homes showed higher epigenetic-BMI (TTP *b*=-0.13, CI=-0.22 to −0.03, *p*=0.009; FFCW-age9: *b*=-0.15, CI=-0.19 to −0.11, *p*<0.001; FFCW-age15: *b*=-0.14, CI=-0.18 to - 0.90, *p*<0.001).

### (5) Children from marginalized race/ethnic groups have higher epigenetic-BMI, reflecting their differing socioeconomic conditions

In TTP and FFCW, African-American/Black and Latinx children had significantly higher epigenetic-BMI compared to White children (**Supplementary Table S8**). Notably, epigenetic-BMI was associated with BMI across all the three socially-constructed race/ethnic groups, although effect sizes varied in subgroup analyses where BMI z-scores were regressed on DNAm-BMI within each group (**Supplementary Table S9**).

Latinx and African-American/Black children tended to be exposed to higher rates of socioeconomic disadvantage compared to White children, and this pattern has previously been observed in these cohorts (Hummer & Hamilton, 2010; Raffington et al., 2022). We therefore tested whether racial/ethnic differences in epigenetic-BMI were accounted for by concurrent socioeconomic disparities. They were: African-American/Black compared to White group differences in epigenetic-BMI were fully statistically accounted for by differences in socioeconomic disadvantage in TTP and FFCW (**Supplementary Table S8**). Latinx compared to White group differences in epigenetic-BMI were fully statistically accounted for by differences in socioeconomic disadvantage in TTP and were substantially reduced in FFCW (though significant differences remained).

### (6) Longitudinal analysis finds that epigenetic-BMI reflects socioeconomic conditions at birth and are not sensitive to observed variation in changes in socioeconomic status across adolescence

In the FFCW, we fit a fixed effect regression model to examine how changes in SES from ages 9 to 15 were associated with changes in epigenetic-BMI from age 9 to 15 years, adjusting for time-invariant variables. Changes in SES and changes in BMI between 9 and 15 were not correlated (*b*=-0.06, CI= −0.15 to 0.04, *p*=0.25; **Supplementary Table S10**). However, SES remained highly stable over time (*b*=0.88, CI= 0.86 to 0.90, *p*<0.001).

Given the stability of SES over child development, our final analysis estimated a random intercept cross-lagged panel model to test whether SES at birth, at age 9, and age 15 predicted epigenetic-BMI at ages 9 and 15. SES at birth predicted epigenetic-BMI at age 9 (b=-0.15, CI=-0.20 to −0.09, p<0.001) (**Supplementary Table S11, Supplemental Figure S3**). As epigenetic-BMI is highly stable from ages 9 to 15 years (b=0.63, CI=0.60 to 0.67, *p*<0.001), this suggests that very early life SES may have a critical influence on lifetime epigenetic scores.

## Discussion

Leveraging twin and longitudinal study designs, we examined (1) whether epigenetic-BMI previously developed in adult blood samples is a valid biomarker of children’s BMI, when measured in saliva DNA-methylation, and (2) whether child epigenetic-BMI is sensitive to social determinants of health in two sociodemographically diverse pediatric cohorts from the US (TTP and FFCW). We found that epigenetic-BMI captured appreciable variance in concurrent BMI z-scores, with effect sizes of a similar magnitude to what has been reported in adults (Hamilton et al., 2019; Wahl et al., 2017): The Δ*R*2 for epigenetic-BMI was 24.3% in TTP, 9.5% in FFCW-age9 and 13.6% in FFCW-age15 after accounting for age, sex, race/ethnicity, pubertal development, and indices of tobacco exposure. Moreover, epigenetic-BMI was higher in children from socioeconomically disadvantaged homes and in children from marginalized racial/ethnic groups, with effect sizes of a similar magnitude to what has been reported for a DNA-methylation predictor of the pace of biological aging (Niccodemi, 2022; Raffington et al., 2022). It remains to be seen whether epigenetic predictors are sensitive to experimental manipulations in socioeconomic resources in realtime, such as cash transfers in early childhood (Troller-Renfree et al., 2022).

Our results validate research that applies epigenetic predictors of aging-related health developed in adult blood samples to pediatric saliva samples. Perhaps counterintuitively, this cross-tissue extension yielded stronger associations than has previously been found using blood samples in pediatric studies. Reed et al. (2020) reported that DNA-methylation measured in blood captured only 1% of the variance in child BMI and 3% in adolescent BMI, and did not prospectively predict future BMI (*see also* Reuben et al., 2020). In contrast, we found that salivary epigenetic-BMI was both a leading and lagging indicator of BMI change. The current results are thus encouraging, as saliva is more feasible to collect than blood in large numbers of participants. In the TTP and FFCWS, for instance, salivary DNA samples are independently collected by participants at home and returned through the mail. Our results contribute to a growing body of evidence that salivary epigenetic profiles of biological aging, physiological decline, health behaviors, and mortality can yield useful biomarkers that are sensitive to social inequalities and are a leading indicator of future health-relevant phenotypes.

Moreover, three of our findings are in line with the developmental origins of health and disease hypothesis and theories of epigenetic regulation of later life health (Barker, 1990; Hayward & Gorman, 2004). First, we observed social stratification of epigenetic-BMI using weights developed in adult samples, which indicates a molecular link between childhood social conditions and adult health. Second, epigenetic-BMI was highly stable across adolescence, which suggests a substantial amount of between-person variation arises earlier in the life course. Third, we found that socioeconomic contexts at birth relative to concurrent socioeconomic contexts in childhood and adolescence best predicted epigenetic-BMI in childhood and adolescence.

There has recently been a paradigm shift in aging research that anchors the onset of damage accumulation to the prenatal period as opposed to later in the life course after the completion of development and the onset of reproductive age (Gladyshev, 2020). Early ontogenetic development is especially sensitive to environmental contexts, given the rapid pace of fetal development and high developmental plasticity. Experiences during this period appear to exert lasting effects on epigenetic predictors of aging-related health in older adults (Schmitz & Duque, 2022). Intriguingly, our twin analyses found that epigenetic-BMI reflected differences in BMI between monozygotic twins. This suggests that salivary epigenetic predictors of health are sensitive to either (1) prenatal and postnatal exposures that differ between monozygotic twins and affect body mass or (2) developmental idiosyncrasy in body mass development.

## Conclusion

Social scientists have long argued that, because adult health and mortality are shaped by childhood conditions, “economic and educational policies that are targeted at children’s well-being are implicitly health policies with effects that reach far into the adult life course” (Hayward & Gorman, 2004). Evaluating the long-term health impact of childhood interventions and policies, however, is challenged by just how long the “long arm of childhood” is: Efforts to improve childhood living conditions could take decades to bear the fruit of reduced mortality and morbidity. The current study, by showing that an epigenetic measure developed in adults is a valid biomarker of childhood BMI, builds on previous work showing that epigenetic predictors can be a molecular “bridge” between childhood and adulthood. It will be valuable for future studies assessing the impact of economic and educational policies in childhood to incorporate DNA-methylation measures as outcomes that are potentially informative about future health payoffs.

## Supporting information

Supplemental Methods Results

Supplemental Results Tables

## Abbreviations

BMI: Body mass index
Epigenetic-BMI: DNA-methylation sites associated with BMI
FFCWS: Future of Families and Child Well-Being Study
SES: Socioeconomic status
TTP: Texas Twin Project

## Article Summary

We show that epigenetic predictors of BMI developed in adults are valid biomarkers of children’s BMI and are sensitive to social inequalities experienced in childhood.

## What’s Known on This Subject

Children who are socioeconomically disadvantaged are at increased risk for high body mass index and multiple diseases in adulthood. Epigenetic mechanisms, including DNA methylation, are thought to be involved in the biological embedding of early life experience that affects lifelong health.

## What This Study Adds

We show that epigenetic predictors of BMI developed in adults are valid biomarkers of children’s BMI and are sensitive to social inequalities experienced in childhood. Our findings are in line with the hypothesis that early life conditions are especially important factors in epigenetic regulation of later life health.

## Contributors Statement

Laurel Raffington and Colter Mitchell developed the study concept and design, performed and supervised data analysis, and drafted the manuscript. Lisa Schneper performed data analysis and provided critical revisions. Travis Mallard, Jonah Fisher, and Liza Vinnik performed data analysis. Kelseanna Hollis-Hansen provided critical revisions. Daniel A. Notterman, Elliot M. Tucker-Drob, and Kathryn P. Harden developed the study concept and design, supervised data analysis, and drafted the manuscript. All authors approved the final manuscript as submitted and agree to be accountable for all aspects of the work.

## References

Almstrup, K., Lindhardt Johansen, M., Busch, A. S., Hagen, C. P., Nielsen, J. E., Petersen, J. H., & Juul, A. (2016). Pubertal development in healthy children is mirrored by DNA methylation patterns in peripheral blood. Scientific Reports, 6(1), 28657. https://doi.org/10.1038/srep28657

Andrews, S. V., Ladd-Acosta, C., Feinberg, A. P., Hansen, K. D., & Fallin, M. D. (2016). “Gap hunting” to characterize clustered probe signals in Illumina methylation array data. Epigenetics & Chromatin, 9(1), 56. https://doi.org/10.1186/s13072-016-0107-z

Aristizabal, M. J., Anreiter, I., Halldorsdottir, T., Odgers, C. L., McDade, T. W., Goldenberg, A., Mostafavi, S., Kobor, M. S., Binder, E. B., Sokolowski, M. B., & O’Donnell, K. J. (2019). Biological embedding of experience: A primer on epigenetics. Proceedings of the National Academy of Sciences, 201820838. https://doi.org/10.1073/pnas.1820838116

Bakulski, K. M., Halladay, A., Hu, V. W., Mill, J., & Fallin, M. D. (2016). Epigenetic Research in Neuropsychiatric Disorders: The “Tissue Issue.” Current Behavioral Neuroscience Reports, 3(3), 264–274. https://doi.org/10.1007/s40473-016-0083-4

Barker, D. J. (1990). The fetal and infant origins of adult disease. BMJ, 301(6761), 1111–1111. https://doi.org/10.1136/bmj.301.6761.1111

Belsky, D. W., Caspi, A., Corcoran, D. L., Sugden, K., Poulton, R., Arseneault, L., Baccarelli, A., Chamarti, K., Gao, X., Hannon, E., Harrington, H. L., Houts, R., Kothari, M., Kwon, D., Mill, J., Schwartz, J., Vokonas, P., Wang, C., Williams, B. S., & Moffitt, T. E. (2022). DunedinPACE, a DNA methylation biomarker of the pace of aging. ELife, 11, e73420. https://doi.org/10.7554/eLife.73420

Biro, F. M., & Wien, M. (2010). Childhood obesity and adult morbidities1–4. 91(5), 1499S–1505S. https://doi.org/10.3945/ajcn.2010.28701B

Braithwaite, D., Moore, D. H., Lustig, R. H., Epel, E. S., Ong, K. K., Rehkopf, D. H., Wang, M. C., Miller, S. M., & Hiatt, R. A. (2009). Socioeconomic status in relation to early menarche among black and white girls. Cancer Causes & Control, 20(5), 713–720. https://doi.org/10.1007/s10552-008-9284-9

Gladyshev, V. N. (2020). The ground zero of organismal life and aging. Trends in Molecular Medicine, S1471491420302173. https://doi.org/10.1016/j.molmed.2020.08.012

Hamaker, E. L., Kuiper, R. M., & Grasman, R. P. P. P. (2015). A critique of the cross-lagged panel model. Psychological Methods, 20(1), 102–116. https://doi.org/10.1037/a0038889

Hamilton, O. K. L., Zhang, Q., McRae, A. F., Walker, R. M., Morris, S. W., Redmond, P., Campbell, A., Murray, A. D., Porteous, D. J., Evans, K. L., McIntosh, A. M., Deary, I. J., & Marioni, R. E. (2019). An epigenetic score for BMI based on DNA methylation correlates with poor physical health and major disease in the Lothian Birth Cohort. International Journal of Obesity, 43(9), 1795–1802. https://doi.org/10.1038/s41366-018-0262-3

Hannon, E., Knox, O., Sugden, K., Burrage, J., Wong, C. C. Y., Belsky, D. W., Corcoran, D. L., Arseneault, L., Moffitt, T. E., Caspi, A., & Mill, J. (2018). Characterizing genetic and environmental influences on variable DNA methylation using monozygotic and dizygotic twins. PLOS Genetics, 14(8), e1007544. https://doi.org/10.1371/journal.pgen.1007544

Harden, K. P., Tucker-Drob, E. M., & Tackett, J. L. (2013). The Texas Twin Project. Twin Research and Human Genetics, 16(01), 385–390. https://doi.org/10.1017/thg.2012.97

Hayward, M. D., & Gorman, B. K. (2004). The long arm of childhood: The influence of early-life social conditions on men’s mortality. Demography, 41(1), 87–107. https://doi.org/10.1353/dem.2004.0005

Hu, D., Xie, F., Xiao, Y., Lu, C., Zhong, J., Huang, D., Wei, J., Jiang, Y., & Zhong, T. (2021). Metformin: A Potential Candidate for Targeting Aging Mechanisms. Aging and Disease, 12(2), 14.

Hummer, R. A., & Hamilton, E. R. (2010). Race and ethnicity in fragile families. The Future of Children, 20(2), 113–131. https://doi.org/10.1353/foc.2010.0003

Joehanes, R., Just, A. C., Marioni, R. E., Pilling, L. C., Reynolds, L. M., Mandaviya, P. R., Guan, W., Xu, T., Elks, C. E., Aslibekyan, S., Moreno-Macias, H., Smith, J. A., Brody, J. A., Dhingra, R., Yousefi, P., Pankow, J. S., Kunze, S., Shah, S. H., McRae, A. F., … London, S. J. (2016). Epigenetic signatures of cigarette smoking. Circulation: Cardiovascular Genetics, 9(5), 436–447. https://doi.org/10.1161/CIRCGENETICS.116.001506

Kankaanpää, A., Tolvanen, A., Heikkinen, A., Kaprio, J., Ollikainen, M., & Sillanpää, E. (2022). The role of adolescent lifestyle habits in biological aging: A prospective twin study. ELife, 11, e80729. https://doi.org/10.7554/eLife.80729

McCartney, D. L., Hillary, R. F., Stevenson, A. J., Ritchie, S. J., Walker, R. M., Zhang, Q., Morris, S. W., Bermingham, M. L., Campbell, A., Murray, A. D., Whalley, H. C., Gale, C. R., Porteous, D. J., Haley, C. S., McRae, A. F., Wray, N. R., Visscher, P. M., McIntosh, A. M., Evans, K. L., … Marioni, R. E. (2018). Epigenetic prediction of complex traits and death. Genome Biology, 19(1), 136. https://doi.org/10.1186/s13059-018-1514-1

Middleton, L. Y. M., Dou, J., Fisher, J., Heiss, J. A., Nguyen, V. K., Just, A. C., Faul, J., Ware, E. B., Mitchell, C., Colacino, J. A., & M. Bakulski, K. (2021). Saliva cell type DNA methylation reference panel for epidemiological studies in children. Epigenetics, 1–17. https://doi.org/10.1080/15592294.2021.1890874

Niccodemi, G. (2022). Pace of aging, family environment and cognitive skills in children and adolescents. 43.

Petersen, A. C., Crockett, L., Richards, M., & Boxer, A. (1988). A self-report measure of pubertal status: Reliability, validity, and initial norms. Journal of Youth and Adolescence, 17(2), 117–133.

Raffington, L., & Belsky, D. W. (2022). Integrating DNA Methylation Measures of Biological Aging into Social Determinants of Health Research. Current Environmental Health Reports, 15.

Raffington, L., Belsky, D. W., Malanchini, M., Tucker-Drob, E. M., & Harden, K. P. (2020). Analysis of socioeconomic disadvantage and pace of aging measured in saliva DNA methylation of children and adolescents. BioRxiv, 2020.06.04.134502. https://doi.org/10.1101/2020.06.04.134502

Raffington, L., Tanksley, P. T., Sabhlok, A., Vinnik, L., Mallard, T., King, L. S., Goosby, B., Harden, K. P., & Tucker-Drob, E. M. (2022). Socially Stratified Epigenetic Profiles Are Associated With Cognitive Functioning in Children and Adolescents. 16.

Reed, Z. E., Suderman, M. J., Relton, C. L., Davis, O. S. P., & Hemani, G. (2020). The association of DNA methylation with body mass index: Distinguishing between predictors and biomarkers. Clinical Epigenetics, 12(1), 50. https://doi.org/10.1186/s13148-020-00841-5

Reuben, A., Sugden, K., Arseneault, L., Corcoran, D. L., Danese, A., Fisher, H. L., Moffitt, T. E., Newbury, J. B., Odgers, C., Prinz, J., Rasmussen, L. J. H., Williams, B., Mill, J., & Caspi, A. (2020). Association of Neighborhood Disadvantage in Childhood With DNA Methylation in Young Adulthood. JAMA Network Open, 3(6), e206095. https://doi.org/10.1001/jamanetworkopen.2020.6095

Sawyer, S. M., Afifi, R. A., Bearinger, L. H., Blakemore, S.-J., Dick, B., Ezeh, A. C., & Patton, G. C. (2012). Adolescence: A foundation for future health. The Lancet, 379(9826), 1630–1640. https://doi.org/10.1016/S0140-6736(12)60072-5

Schmitz, L. L., & Duque, V. (2022). In utero exposure to the Great Depression is reflected in late-life epigenetic aging signatures. Proceedings of the National Academy of Sciences, 119(46), e2208530119. https://doi.org/10.1073/pnas.2208530119

Shah, S., Bonder, M. J., Marioni, R. E., Zhu, Z., McRae, A. F., Zhernakova, A., Harris, S. E., Liewald, D., Henders, A. K., Mendelson, M. M., Liu, C., Joehanes, R., Liang, L., Levy, D., Martin, N. G., Starr, J. M., Wijmenga, C., Wray, N. R., Yang, J., … van Zwet, E. W. (2015). Improving phenotypic prediction by combining genetic and epigenetic associations. The American Journal of Human Genetics, 97(1), 75–85. https://doi.org/10.1016/j.ajhg.2015.05.014

St-Onge, M.-P., & Gallagher, D. (2010). Body composition changes with aging: The cause or the result of alterations in metabolic rate and macronutrient oxidation? Nutrition, 26(2), 152–155. https://doi.org/10.1016/j.nut.2009.07.004

Sumner, J. A., Colich, N. L., Uddin, M., Armstrong, D., & McLaughlin, K. A. (2019). Early experiences of threat, but not deprivation, are associated with accelerated biological aging in children and adolescents. Biological Psychiatry, 85(3), 268–278. https://doi.org/10.1016/j.biopsych.2018.09.008

Trejo Banos, D., McCartney, D. L., Patxot, M., Anchieri, L., Battram, T., Christiansen, C., Costeira, R., Walker, R. M., Morris, S. W., Campbell, A., Zhang, Q., Porteous, D. J., McRae, A. F., Wray, N. R., Visscher, P. M., Haley, C. S., Evans, K. L., Deary, I. J., McIntosh, A. M., … Robinson, M. R. (2020). Bayesian reassessment of the epigenetic architecture of complex traits. Nature Communications, 11(1), Article 1. https://doi.org/10.1038/s41467-020-16520-1

Troller-Renfree, S. V., Costanzo, M. A., Duncan, G. J., Magnuson, K., Gennetian, L. A., Yoshikawa, H., Halpern-Meekin, S., Fox, N. A., & Noble, K. G. (2022). The impact of a poverty reduction intervention on infant brain activity. Proceedings of the National Academy of Sciences, 119(5). https://doi.org/10.1073/pnas.2115649119

US Department of Health and Human Services (USDHHS). (2012). (Preventing Tobacco Use Among Youth and Young Adults: A Report of the Surgeon General USDHHS, Centers for Disease Control and Prevention, National Center for Chronic Disease Prevention and Health Promotion, Office on Smoking and Health).

Wahl, S., Drong, A., Lehne, B., Loh, M., Scott, W. R., Kunze, S., Tsai, P.-C., Ried, J. S., Zhang, W., Yang, Y., Tan, S., Fiorito, G., Franke, L., Guarrera, S., Kasela, S., Kriebel, J., Richmond, R. C., Adamo, M., Afzal, U., … Chambers, J. C. (2017). Epigenome-wide association study of body mass index, and the adverse outcomes of adiposity. Nature, 541(7635), 81–86. https://doi.org/10.1038/nature20784

Watowich, M. M. (2022). Natural disaster and immunological aging in a nonhuman primate. Proceedings of the National Academy of Sciences, 119(8). https://doi.org/e2121663119

