## Supplemental Methods Results for "Measuring the long arm of childhood in real-time: Epigenetic predictors of BMI and social determinants of health across childhood and adolescence"

#### DNA-methylation

*DNA-methylation preprocessing Texas Twins.* Saliva samples were collected during a laboratory visit using Oragene kits (DNA Genotek, Ottawa, ON, Canada). DNA extraction and methylation profiling was conducted by Edinburgh Clinical Research Facility (UK). The Infinium MethylationEPIC BeadChip kit (Illumina, Inc., San Diego, CA) was used to assess methylation levels at 850,000 methylation sites.

DNA-methylation preprocessing was primarily conducted with the 'minfi' package in R (Aryee et al., 2014). Within-array normalization was performed to address array background correction, red/green dye bias, and probe type I/II correction, and it has been noted that at least part of the probe type bias is a combination of the first two factors (Dedeurwaerder et al., 2014). Noob preprocessing as implemented by minfi's "preprocessNoob" (Triche et al., 2013) is a background correction and dye-bias equalization method that has similar within-array normalization effects on the data as probe type correction methods such as BMIQ (Teschendorff et al., 2013).

CpG probes with detection $p > 0.01$ and fewer than 3 beads in more than 1% of the samples and probes in cross-reactive regions were excluded (Pidsley et al., 2016). None of these failed probes overlapped with the probes used for epigenetic predictors. 44 samples were excluded because (1) they showed low intensity probes as indicated by the log of average methylation <9 and their detection p was > 0.01 in >10% of their probes, (2) their self-reported and methylation-estimated sex mismatch, and/or (3) their self-reported and DNA-estimated sex mismatch. Cell composition of immune and epithelial cell types (*i.e.,* CD4+ T-cell, natural killer cells, neutrophilseosinophils,

B cells, monocytes, CD8+ T-cell, and granulocytes) were estimated using a newly developed child saliva reference panel implemented in the *R* package "BeadSorted.Saliva.EPIC" within "ewastools" (Middleton et al., 2021). Surrogate variable analysis was used to correct methylation values for batch effects using the "combat" function in the SVA package (Johnson et al., 2007).

*DNA-methylation preprocessing FFCW.* DNA extraction and methylation profiling for FFCW was conducted by the Notterman Lab of Princeton University and the Pennsylvania State University College of Medicine Genome Sciences Center. Due to the timing of assay completion 40% of the FFCW saliva samples were completed using the Illumina 450K chip and the remaining 60% used the Illumina EPIC chip. Methods for the two chips were standardized as much as possible, but all analyses were run separately for 450 and EPIC and then meta-analyzed. *450K* DNA-methylation image data were processed in R statistical software (4.1) using the ENmix package (Xu et al., 2016). The red and green image pairs ($n_{samples}$ =1811) were read into *R* and the ENmix *preprocessENmix* and *rcp* functions were used to normalize dye bias, apply background correction, and adjust for probe-type bias. The majority of sample filtering was applied using the ewastools packages (Just & Heiss, 2018). Samples were excluded using the following criteria: if >10% of DNA-methylation sites had detection p-value >0.01 ($n_{samples}$ =34), if there was sex discordance between DNA-methylation predicted sex and recorded sex ($n_{samples}$ =11), or if two sequential samples from the same individual exhibited genetic discordance between visits ($n_{samples}$ =27). ENmix *QCinfo* function identified samples with outlier methylation values which were cut ($n_{samples}$ =6). Technical replicates were removed $n_{samples}$ =49). This gave us our final analytic sample (n=1684). DNA-methylation sites were removed if they had detection p-value >0.01 in 5% of samples (n=33,376). Relative proportions of immune and epithelial cell types were estimated from DNA-methylation measures using a childhood saliva reference panel (Middleton et al., 2021).

*EPIC* DNA-methylation image data were processed in R statistical software (4.1) using the ENmix package (Xu et al., 2016).

The red and green image pairs ($n_{samples}$ =2558) were read into *R* and the ENmix *preprocessENmix* and *rcp* functions were used to normalize dye bias, apply background correction, and adjust for probe-type bias. The majority of sample filtering was applied using the ewastools packages (Just & Heiss, 2018). We dropped samples using the following criteria: if >10% of DNA-methylation sites had detection p-value >0.05 ($n_{samples}$ =63), if there was sex discordance between DNA-methylation predicted sex and recorded sex (n=12), or if two sequential samples from the same individual exhibited genetic discordance between visits (n=30). ENmix *QCinfo* function identified samples with outlier methylation values which were cut (n=1) or samples that failed bisulfite conversion ($n_{samples}$ =7). Technical replicates were removed (n=168). This gave us our final analytic sample ($n_{samples}$ =2277). DNA-methylation sites were removed if they had detection p-value >0.05 in 5% of samples (n=127,275). Relative proportions of immune and epithelial cell types were estimated from DNA-methylation measures using a childhood saliva reference panel (Middleton et al., 2021).

**Genetics**

*Texas Twins genotyping, imputation, and preprocessing*. DNA samples were genotyped at the University of Edinburgh using the Illumina Infinium PsychArray, which assays ~590,000 single nucleotide polymorphisms (SNPs), insertions-deletions (indels), copy number variants (CNVs), structural variants, and germline variants across the genome. Genetic data was subjected to quality control procedures recommended for chip-based genomic data (Anderson et al., 2010; Turner et al., 2011). Briefly, samples were excluded on the basis of poor call rate (< 98%) or inconsistent self-reported and biological sex, while variants were excluded if missingness exceeded 2%. As further variant-level filtering has been shown to have a detrimental effect on imputation quality (Roshyara et al., 2014), quality control thresholds for minor allele frequency (MAF) and Hardy–Weinberg equilibrium (HWE) were applied after phasing and imputation. Untyped markers were imputed on the Michigan Imputation Server (https://imputationserver.sph.umich.edu). Specifically, genotypes were phased and imputed with Eagle v2.4 and Minimac4 (v1.5.7), respectively, while using the 1000 Genomes Phase 3 v5 reference panel (Auton et al., 2015). To ensure that only high-quality typed and imputed markers were used for analysis, variants were excluded if they had a MAF < 1e-3, a HWE p-value < 1e-6, or an imputation quality score < .90. These procedures produced a final set of 4,703,309 genetic markers to be used in analyses.

*FFCW* Specimen processing was conducted at the Notterman laboratory at Princeton University and the Pennsylvania State University College of Medicine Genome Sciences Center from 2015-2019 (R01-HD-36916, R01-HD-39135, R01-HD-40421). Genotype data on FFCWS participants was obtained using the Illumina PsychChip_v1-1 and PsychChip_15048346_B. Individuals with missing call rates >2%, SNPs with missing call rates >2%, and chromosomal anomalies were removed. 3,074 individuals and 273,800 SNPs passed filters and QC. PC analysis was performed to identify analytic genomic group outliers and to provide sample eigenvectors as covariates in the statistical model used for association testing to adjust for possible population stratification. SNPs used for PC analysis were selected by linkage disequilibrium (LD) pruning from an initial pool consisting of all autosomal SNPs with a missing call rate < 2% and minor allele frequency (MAF) > 5%, and excluding any SNPs with a discordance between HapMap controls genotyped along with the study samples and those in the external HapMap data set. In addition, we excluded the HLA, 8p23, and 17q21.31 regions from the initial pool. We categorized participants through PC analysis using the aforementioned filtering criteria into analytic genomic groups based on genome-wide SNP similarity to genomic reference groups (commonly referred to using geographical labels as European, n=475; Admixed African, n=1640; Admixed Hispanic, n=959 groups). Then, PC analysis was run again within each group to create sample eigenvectors for covariates in the statistical model used for association testing to adjust for possible population stratification within each analytic group (*i.e.,* within-analytic-group PCs).

*Polygenic scores of BMI.*

Genetic data was used to calculate a polygenic score of BMI (PGS-BMI). The PGS-BMI is an approximate indicator of an individual's genetic liability for developing high levels of BMI. We computed polygenic scores of BMI on the basis of a meta-analysis of genome-wide association studies for body fat distribution in 694 649 individuals of high genetic similarity to European reference groups (Pulit et al., 2019). Polygenic scores were residualized for the top 10 principal components of genetic similarity to reference groups as in FFCW. PGS-BMI analyses were restricted to individuals of high genomic similarity to European and "Ad Mixed American" reference groups in order to reduce the risk of spurious findings due to population stratification (see https://useast.ensembl.org/Help/Faq?id=532).

### Supplemental Results

#### Cross-sectional analyses show that salivary epigenetic-BMI provides complementary information to measured genetic variants

Previous genome-wide association studies of adult BMI have estimated correlations with measured DNA sequence variants. This information can be applied in new samples to compute polygenic scores, which represent an individual's aggregate genetic liability toward developing high BMI (PGS-BMI). PGSs have limitations: They also capture environmental processes such as uncorrected population stratification, "genetic nurture," and gene-environment correlations, and they do not capture the effects of all genetic variation, including rare variants (Raffington et al., 2020; Young et al., 2019). Nonetheless, in previous work in adults, epigenetic-BMI and PGS-BMI provided complementary prediction with respect to BMI (Shah et al., 2015; Trejo Banos et al., 2020). One previous pediatric study of epigenetic-BMI in blood also found that epigenetic-BMI and PGS-BMI measures captured largely independent variation in child and adolescent BMI (Reed et al., 2020).

Due to different patterns of linkage disequilibrium across ancestries, and possible gene × environment interactions, it is expected that PGS will have imperfect portability to groups with disparate ancestries relative to the discovery sample (Martin et al., 2019). Consequently, we report analyses separately by DNA-based ancestry estimates (Supplemental Table S6). We regressed BMI z-scores on PGS-BMI residualized for principal components of genetic similarity to reference groups (see Methods). In both cohorts and all ancestral groups, children with higher PGS-BMI had higher BMI (see **Supplementary Table S6, Supplementary Figure S1**).

Next, we tested whether epigenetic-BMI remained associated with childhood BMI after accounting for PGS-BMI and it did (see **Supplementary Table S6)**. Consistent with results from previous studies (Reed et al., 2020; Shah et al., 2015; Trejo Banos et al., 2020), the variation in BMI explained by epigenetic-BMI and PGS-BMI was largely additive (see **Supplementary Figure S1).**

**Supplemental Figures**

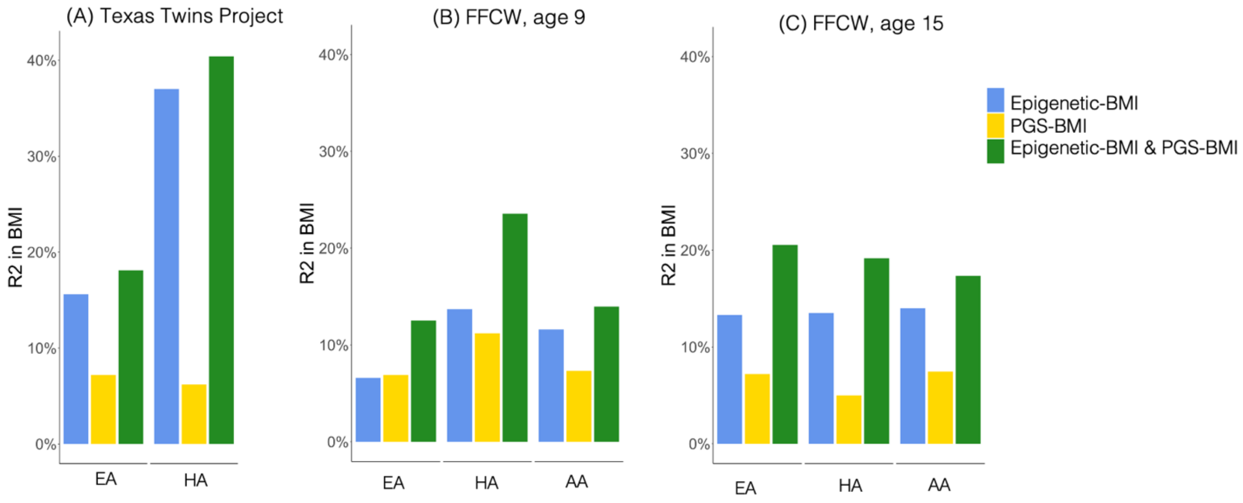

**Figure S1. Measures of epigenetic-BMI and polygenic scores (PGS-BMI) uniquely contribute to prediction of BMI.** Participants were categorized into genetic analytic groups based on similarity to genomic reference groups (EA, HA, AA) due to risk of social stratification confounding in PGS analyses (see Supplemental Methods). Results presented for 8- to 18-year-old children from the Texas Twin Project (TTP), and in 9-year-old children and 15-year-old children from Future of Families and Child Wellbeing Study (FFCWS).

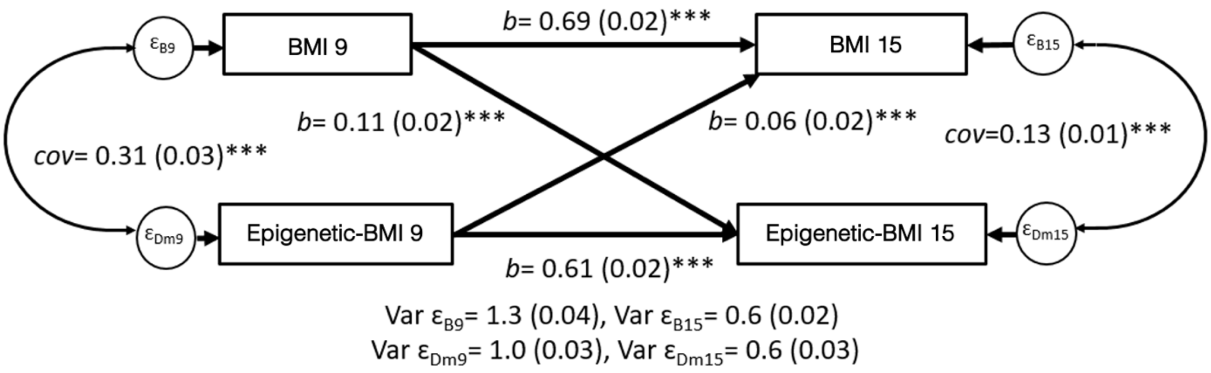

**Figure S2. Cross Lagged Model of epigenetic-BMI and BMI in FFCWS.** BMI is the CDC z-score for BMI for age. Epigenetic-BMI was residualized for cell distributions and technical artifacts and normalized with a mean=0, SD=1. Coefficients not shown are controls that influence age 9 BMI and age 9 epigenetic-BMI, including: self-reported race, sex assigned at birth, SES at birth, if mom smoked while pregnant (see Supplement Table S6). Standard deviations of the Residual error variances of BMI and epigenetic-BMI are in parenthesis after their variance estimate. *p<0.05, **p<0.01,***p<0.001

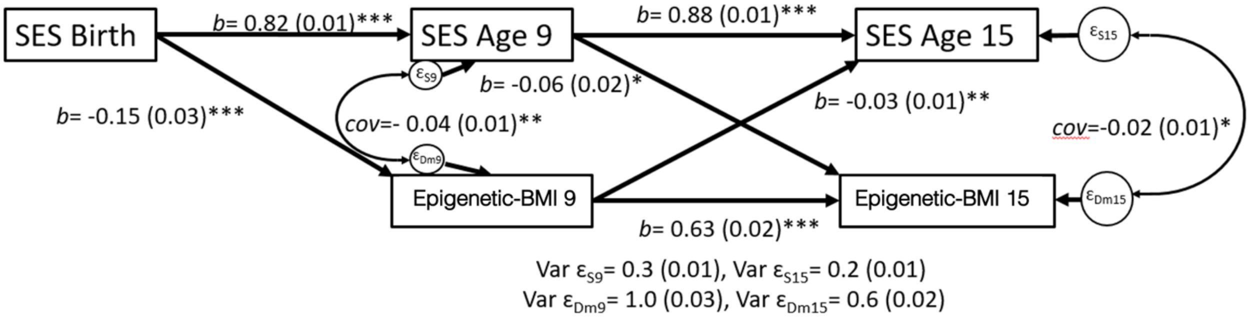

**Figure S3. Cross Lagged Model of SES and epigenetic-BMI Profiles in FFCWS.** Epigenetic-BMI was residualized for cell distributions and technical artifacts. SES and epigenetic-BMI are normalized with a mean=0, SD=1. Coefficients not shown are controls that influence age 9 SES and Age 9 epigenetic-BMI, including: self-reported race, sex assigned at birth, and if mom smoked while pregnant (see Supplemental Table 10). Standard deviations of the residual error variances of SES and epigenetic-BMI are in parenthesis after their variance estimate. *p<0.05, **p<0.01,***p<0.001

### Supplemental References

Anderson, C. A., Pettersson, F. H., Clarke, G. M., Cardon, L. R., Morris, A. P., & Zondervan, K. T. (2010). Data quality control in genetic case-control association studies. *Nature Protocols*, *5*(9), 1564–1573. https://doi.org/10.1038/nprot.2010.116

Aryee, M. J., Jaffe, A. E., Corrada-Bravo, H., Ladd-Acosta, C., Feinberg, A. P., Hansen, K. D., & Irizarry, R. A. (2014). Minfi: A flexible and comprehensive Bioconductor package for the analysis of Infinium DNA methylation microarrays. *Bioinformatics*. https://doi.org/10.1093/bioinformatics/btu049

Auton, A., Abecasis, G. R., Altshuler, D. M., Durbin, R. M., Abecasis, G. R., Bentley, D. R., Chakravarti, A., Clark, A. G., Donnelly, P., Eichler, E. E., Flicek, P., Gabriel, S. B., Gibbs, R. A., Green, E. D., Hurles, M. E., Knoppers, B. M., Korbel, J. O., Lander, E. S., Lee, C., … National Eye

Institute, N. (2015). A global reference for human genetic variation. *Nature*, *526*(7571), Article 7571. https://doi.org/10.1038/nature15393

Johnson, W. E., Li, C., & Rabinovic, A. (2007). Adjusting batch effects in microarray expression data using empirical Bayes methods. *Biostatistics*, *8*(1), 118–127. https://doi.org/10.1093/biostatistics/kxj037

Just, A. C., & Heiss, J. A. (2018). *ewastools: EWAS Tools*. R package version 16.

Martin, A. R., Kanai, M., Kamatani, Y., Okada, Y., Neale, B. M., & Daly, M. J. (2019). Clinical use of current polygenic risk scores may exacerbate health disparities. *Nature Genetics*, *51*(4), 584–591. https://doi.org/10.1038/s41588-019-0379-x

Middleton, L. Y. M., Dou, J., Fisher, J., Heiss, J. A., Nguyen, V. K., Just, A. C., Faul, J., Ware, E. B., Mitchell, C., Colacino, J. A., & M. Bakulski, K. (2021). Saliva cell type DNA methylation reference panel for epidemiological studies in children. *Epigenetics*, 1–17. https://doi.org/10.1080/15592294.2021.1890874

Pidsley, R., Zotenko, E., Peters, T. J., Lawrence, M. G., Risbridger, G. P., Molloy, P., Van Djik, S., Muhlhausler, B., Stirzaker, C., & Clark, S. J. (2016). Critical evaluation of the Illumina MethylationEPIC BeadChip microarray for whole-genome DNA methylation profiling. *Genome Biology*, *17*(1), 208. https://doi.org/10.1186/s13059-016-1066-1

Pulit, S. L., Stoneman, C., Morris, A. P., Wood, A. R., Glastonbury, C. A., Tyrrell, J., Yengo, L., Ferreira, T., Marouli, E., Ji, Y., Yang, J., Jones, S., Beaumont, R., Croteau-Chonka, D. C., Winkler, T. W., Hattersley, A. T., Loos, R. J. F., Hirschhorn, J. N., Visscher, P. M., … Lindgren, C. M. (2019). Meta-analysis of genome-wide association studies for body fat distribution in 694 649 individuals of European ancestry. *Human Molecular Genetics*, *28*(1), 166–174. https://doi.org/10.1093/hmg/ddy327

Raffington, L., Mallard, T., & Harden, K. P. (2020). Polygenic scores indevelopmental psychology: Invite genetics In, leave biodeterminism behind. *Annual Review of Developmental Psychology*, *2*(1), null. https://doi.org/10.1146/annurev-devpsych-051820-123945

Reed, Z. E., Suderman, M. J., Relton, C. L., Davis, O. S. P., & Hemani, G. (2020). The association of DNA methylation with body mass index: Distinguishing between predictors and biomarkers. *Clinical Epigenetics*, *12*(1), 50. https://doi.org/10.1186/s13148-020-00841-5

Roshyara, N. R., Kirsten, H., Horn, K., Ahnert, P., & Scholz, M. (2014). Impact of pre-imputation SNP-filtering on genotype imputation results. *BMC Genetics*, *15*(1), 88. https://doi.org/10.1186/s12863-014-0088-5

Shah, S., Bonder, M. J., Marioni, R. E., Zhu, Z., McRae, A. F., Zhernakova, A., Harris, S. E., Liewald, D., Henders, A. K., Mendelson, M. M., Liu, C., Joehanes, R., Liang, L., Levy, D., Martin, N. G., Starr, J. M., Wijmenga, C., Wray, N. R., Yang, J., … van Zwet, E. W. (2015). Improving phenotypic prediction by combining genetic and epigenetic associations. *The American Journal of Human Genetics*, *97*(1), 75–85. https://doi.org/10.1016/j.ajhg.2015.05.014

Trejo Banos, D., McCartney, D. L., Patxot, M., Anchieri, L., Battram, T., Christiansen, C., Costeira, R., Walker, R. M., Morris, S. W., Campbell, A., Zhang, Q., Porteous, D. J., McRae, A. F., Wray, N. R., Visscher, P. M., Haley, C. S., Evans, K. L., Deary, I. J., McIntosh, A. M., … Robinson, M. R. (2020). Bayesian reassessment of the epigenetic architecture of complex traits. *Nature Communications*, *11*(1), Article 1. https://doi.org/10.1038/s41467-020-16520-1

Triche, T. J., Weisenberger, D. J., Van Den Berg, D., Laird, P. W., & Siegmund, K. D. (2013). Low-level processing of Illumina Infinium DNA Methylation BeadArrays. *Nucleic Acids Research*, *41*(7), e90–e90. https://doi.org/10.1093/nar/gkt090

Turner, S., Armstrong, L. L., Bradford, Y., Carlson, C. S., Crawford, D. C., Crenshaw, A. T., de Andrade, M., Doheny, K. F., Haines, J. L., Hayes, G., Jarvik, G., Jiang, L., Kullo, I. J., Li, R., Ling, H., Manolio, T. A., Matsumoto, M., McCarty, C. A., McDavid, A. N., … Ritchie, M. D. (2011). Quality Control Procedures for Genome-Wide Association Studies. *Current Protocols in Human Genetics*, *68*(1), 1.19.1-1.19.18. https://doi.org/10.1002/0471142905.hg0119s68

Xu, Z., Niu, L., Li, L., & Taylor, J. A. (2016). ENmix: A novel background correction method for Illumina HumanMethylation450 BeadChip. *Nucleic Acids Research*, *44*(3), e20–e20. https://doi.org/10.1093/nar/gkv907

Young, A. I., Benonisdottir, S., Przeworski, M., & Kong, A. (2019). Deconstructing the sources of genotype-phenotype associations in humans. *Science*, *365*(6460), 1396–1400. https://doi.org/10.1126/science.aax3710
